# Accounting for recurrent mutation in the frequency spectrum of rare alleles

**DOI:** 10.64898/2026.05.29.728884

**Authors:** Deepjyoti Ghosh, Kyle Williams, Joshua Schraiber, Yuval Simons

## Abstract

As whole-genome and whole-exome datasets increase in size, they uncover alleles at lower and lower frequencies in the population. Samples of rare alleles often include recurrent mutations, where derived alleles are identical by state and not by descent. As a result, the site frequency spectrum (SFS) becomes challenging to analyze because it is strongly dependent on the mutation rate. To overcome this hurdle, we define the single mutation frequency spectrum (SMFS), which is the frequency spectrum of alleles descendant from a single mutational event. For rare alleles, the SFS with recurrent mutation is then a weighted sum of the convolutions of the SMFS with itself. This simple, yet powerful, model decouples recurrent mutation from the population genetic processes giving rise to the SMFS, such as genetic drift and selection. We show how both forward-in-time and backward-in-time models with recurrent mutations can be recast in terms of the SMFS. We then develop a method for combinatorial hierarchic estimation of the SMFS (which we name CHES). We apply this simple, yet robust, method to a human exome sequencing dataset to show that the SMFS with recurrent mutation can account for SFS differences between low and high mutation rate sites. The inferred SMFS shows an approximate scaling law with allele frequencies inconsistent with both a constant population size and an exponentially growing population model. Lastly, we use our model to compare the expected and observed proportions of missense and stop-gain mutations in the human exome, using this disparity to infer the strength of selection on these classes of mutations. Our combined results show how the SMFS can explain the dependency of the SFS on the mutation rate and how it reflects human demographic and evolutionary history.

## 1 Introduction

As whole-genome and whole-exome sequencing datasets in humans scale up to include hundreds of thousands of samples, it has become feasible to detect mutations with increasingly lower allele frequencies^1–6^. When multiple copies of these rare alleles are observed in such large datasets, they often represent recurrent mutations — instances where derived alleles are identical by state (IBS) rather than by descent (IBD)^2,5,7–9^. This recurrence of mutations complicates the analysis of the site frequency spectrum (SFS)^7–10^, which is the distribution of allele frequencies in a sampled population.

The SFS is central to many statistical and population genetic methods for inferring demographic history and selection from genetic data^11–15^. However, many such methods and their underlying models explicitly or implicitly assume no recurrent mutation^13,16–20^ (an “infinite sites” approximation^21^). This approximation works well when sample sizes are not overly large, but breaks down when applied to large-scale genomic datasets from projects like the UK Biobank^6^ and gnomAD^5^. At these extremely large sample sizes, sampling of rare variants arising from recurrent mutations becomes more common, rendering the SFS strongly dependent on the mutation rate and thus difficult to analyze^7,8,22^. For example, a dataset including one million human non-Finnish European haplotypes exhibited a pronounced difference between the SFS of CpG and non-CpG transition driven mainly by differences in mutation rate^7,8^ (see Fig. 1).

**Figure 1.**
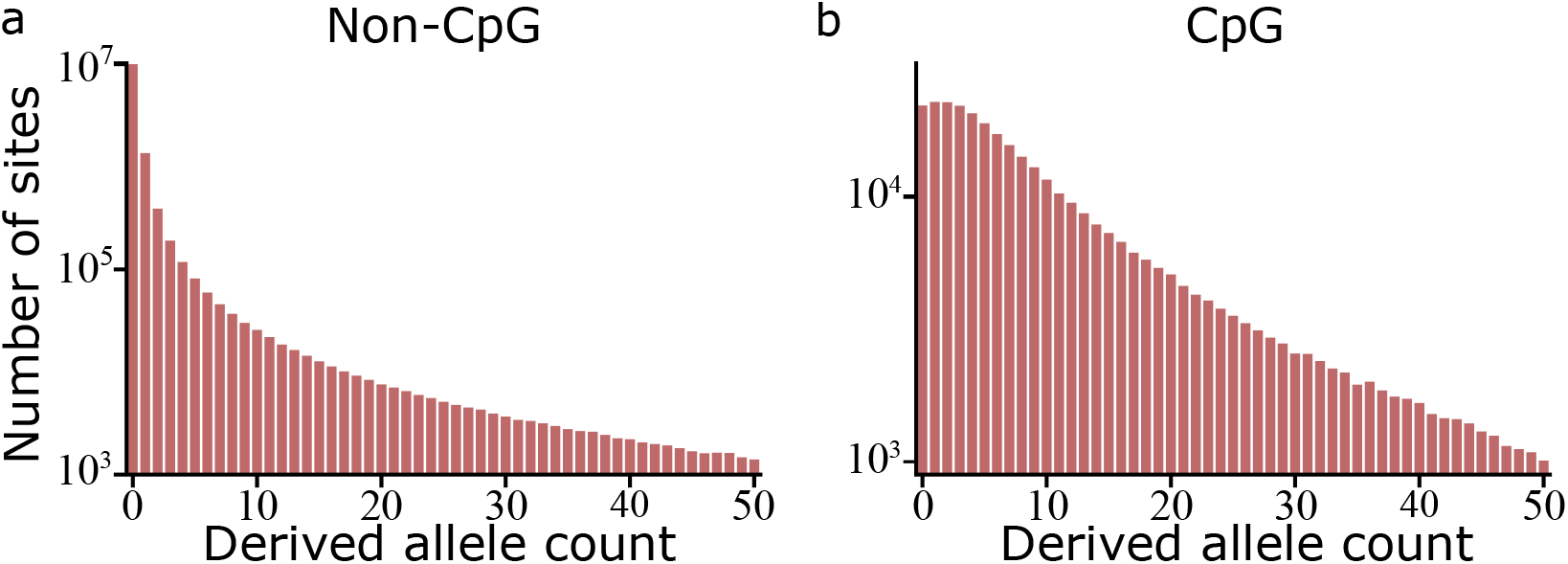
Site frequency spectrum for synonymous mutations in a whole exome dataset with 1,000,000 haplotypes (Schraiber et al.^36^). For sites with multiple derived alleles, we consider the different alleles separately. We see a marked difference in the shape of the SFS between (a) regular mutations and (b) C to T mutations at methylated CpG sites (which have elevated mutation rates).

The distribution of rare variants in these biobank-scale genomic datasets provides the basis for inference of variation in human mutation rate, of recent human demographic history, and of the strength of natural selection^5,13–15,23–28^. Yet this inference is hampered due to a lack of models of the SFS that account for the possibility of recurrent mutation, i.e., for the possibility that multiple copies of an allele may have arisen independently rather than from a common ancestral mutation^10^. While some theoretical works have addressed the challenge of incorporating recurrent mutation into the analysis of the SFS^7,8,22,29–32^, most of them do not specifically consider rare variation. This limits their usefulness in modeling human populations—since humans have a low mutation rate, recurrent mutations necessarily occur at low frequencies.

However, the low mutation rate may also simplify the problem of analyzing the SFS under recurrent mutation. At low allele frequencies, the frequency dynamics of alleles descendant from different mutations at the same site are independent of each other^33^, and at low per generation mutation rate these dynamics are also independent of the mutation rate^8^. This suggests the frequency spectrum of alleles identical-by-descent from a single mutational event would conform to an “infinite sites” like approximation, with a frequency distribution whose shape is independent of the mutation rate. Only when recurrent mutations are indistinguishable does the observed frequency spectrum become dependent on the mutation rate—as the sum of alleles descendant from different mutational events, it depends directly on the rate at which such events occur. That is, it is not the evolutionary dynamics that give rise to mutation rate dependence, but the fact that we can’t distinguish identical mutations.

This description of the SFS with recurrent mutation suggests a simple approach to modeling the SFS as a compound Poisson process. We define the frequency spectrum of segregating alleles IBD from a single mutational event as the Single Mutation Frequency Spectrum (SMFS). The observed SFS with recurrent mutation is then a convolution of the SMFS that depends on the number of mutational events sampled, which is generally unknown (but see strategies to estimate it in the Discussion below). As is standard under low mutation rates, we assume that the number of mutational events is a Poisson process with mean proportional to the mutation rate.

We show that this model unifies and simplifies the results of two recent approaches to accounting for recurrent mutations in the SFS: A forward-in-time approach^7^ modeling the dynamics of rare alleles as a branching process using sums over Bell’s polynomials of transition matrices to account for recurrent mutations; and a backward-in-time coalescent approach^8^ that relates the SFS with recurrent mutations to a combinatorial sum over summaries of the gene genealogy. While mathematically equivalent to both of these previous approaches, and very similar to the Poisson Random field approach^11,29,34,35^, our model separates the effects of recurrent mutation from those of other population genetic processes.

This separation makes our approach mathematically and computationally simple and allows us to develop a combinatorial hierarchical method, which we call CHES, to estimate the SMFS from the SFS. The estimated SMFS then provides a mutation rate independent signal of the underlying population genetic processes. We show how the SMFS in a large human dataset shows evidence of recent explosive population growth, and how the SFS of missense and stop-gain coding mutations deviates from the SMFS predictions, allowing us to estimate the fraction of such mutations under strong selection.

## 2 Model

### 2.1 Intuition from the infinite sites model

Many population genetic models employ what is known as an infinite sites model (ISM)^21^. The ISM assumes that there is an effectively infinite number of sites where a mutation can occur, and therefore every new mutation occurs at a different site, i.e., there are no recurrent mutations. This is an appropriate assumption when the mutation rate is low enough for recurrent mutations to be exceedingly rare. Under the ISM, the derived alleles at a segregating site are always identical by descent, i.e., they are derived from a single mutational event.

Under the ISM, the probability of a site segregating in a sample is proportional to the mutation rate, *u*, and the population size, *N*_*e*_, that is, *P*(*segregating*) = 2 ·*N*_*e*_ ·*u* ·*ξ* for population size *N*_*e*_, with some constant *ξ*, determined by population genetic factors such as selection and demographic history^33^. (Note that when the population size isn’t constant there is a degree of arbitrariness in choice of the reference population size *N*_*e*_, and the choice of *N*_*e*_ rescales *ξ*.) The SFS (including non-segregating alleles) is then:

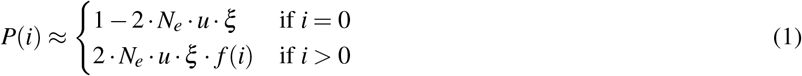

with *f* (*i*) being the mutation rate independent SFS of segregating alleles, with 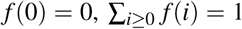. This is a slight rephrasing of the classical notation which will prove useful in our analysis below. Under neutral Wright-Fisher dynamics with panmixia and a constant population size, 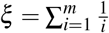 and 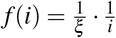 with *m* being the sample size^37^.

Inspired by the ISM, we developed an analogous model to describe a different scenario, when rare derived alleles at a site may arise from more than one mutational event. We refer to this as an “infinite ancestral background model”: the population has infinitely many copies of the ancestral allele. Each new mutation mutates one of the ancestral alleles to a derived allele, where we assume we cannot differentiate the different derived alleles. Under this model, different derived alleles may be identical-by-state and not by descent. Because the population is infinite, the evolutionary fate of the allele introduced by each mutational event is independent of the fate of alleles descendant from other mutational events. We refer to the frequency distribution of alleles descendant from exactly one mutational event, *f* (*i*), as the Single Mutation Frequency Spectrum (SMFS). The SMFS would be identical to *f* (*i*) in our above description of the infinite sites frequency spectrum for a low enough mutation rate.

We call this an “infinite ancestral background model” to emphasize its difference from the ISM—we take the population size, not the number of sites, to infinity, thereby allowing multiple mutations per site, all on the same ancestral background and all with the same frequency spectrum, the SMFS. This is mathematically equivalent to an infinite sites model in which we sum allele counts over many sites such that the sum of their 2 ·*N*_*e*_ ·*u* ·*ξ* may be greater than one. This is a generalization of the model introduced by Desai and Plotkin^29^. We will now make this simple model mathematically explicit.

### 2.2 Mathematical description

Let *f* (*i*) denote the SMFS, with *i* the count of alleles IBD from a single mutation. *f* (*i*) is the probability distribution of the number of sampled alleles descendant from a single mutational event. Note that by definition, *f* (0) = 0 and the SMFS is normalized to 1, i.e., ∑_*i* ≥ 0_ *f* (*i*) = 1.

Derived alleles at a single site may be descendant from one, two, or any number of mutational events. If, and only if, the number of mutational events sampled is 0 can we observe no derived alleles. If the number of mutational events sampled is one, then the distribution of the number of derived alleles is simply *f* (*i*). If *k* mutational events are sampled, then the number of alleles descendant from each mutational event has the distribution *f* (*i*), and the distribution of the overall number of derived alleles is *f* ^\**k*^(*i*), where *f* ^\**k*^(*i*) is the *k*-th convolution of *f* with itself and is defined by:

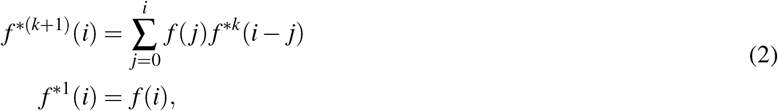

where,

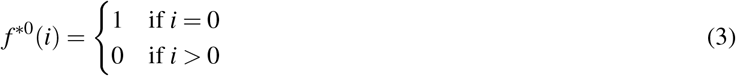

for the distribution when no mutational events are sampled.

Note that since each sampled mutational event contributes at least one derived allele then *f* ^\**k*^(*i*) = 0 for all *i* < *k*. We can prove this by induction. For *k* = 1, *f* ^*1^(0) = *f* (0) = 0. If we assume it is true for *k*, then for *i* < *k* + 1:

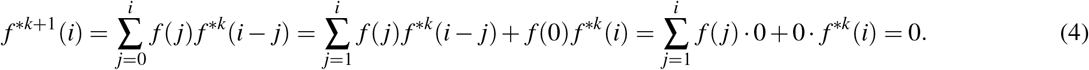

This truncation will later become the foundation of our hierarchical inference procedure.

We model the number of mutational events sampled as a Poisson process with mean *λ* = 2 · *N*_*e*_ · *u* · *ξ*,

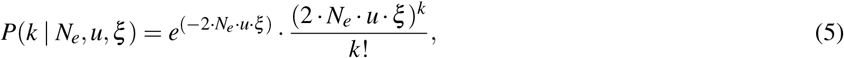

where, as before, *N*_*e*_ is the effective population size, *u* is the mutation rate, and *ξ* is a sample size dependent constant representing the number of sampled mutational events per mutational input. We can combine these results to obtain an expression for the SFS. That is, the probability of observing an allele count of *i* is:

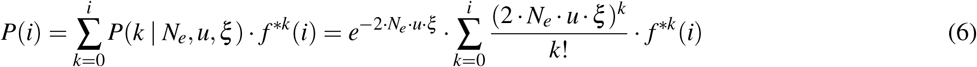

where the sum runs only up to *i* because for *k* > *i, f* ^\**k*^(*i*) = 0. As expected, the SFS is mutation rate dependent. As we can see in Fig. 2a, for low *u* (when 2 · *N*_*e*_ · *u* · *ξ* ≪ 1):

**Figure 2.**
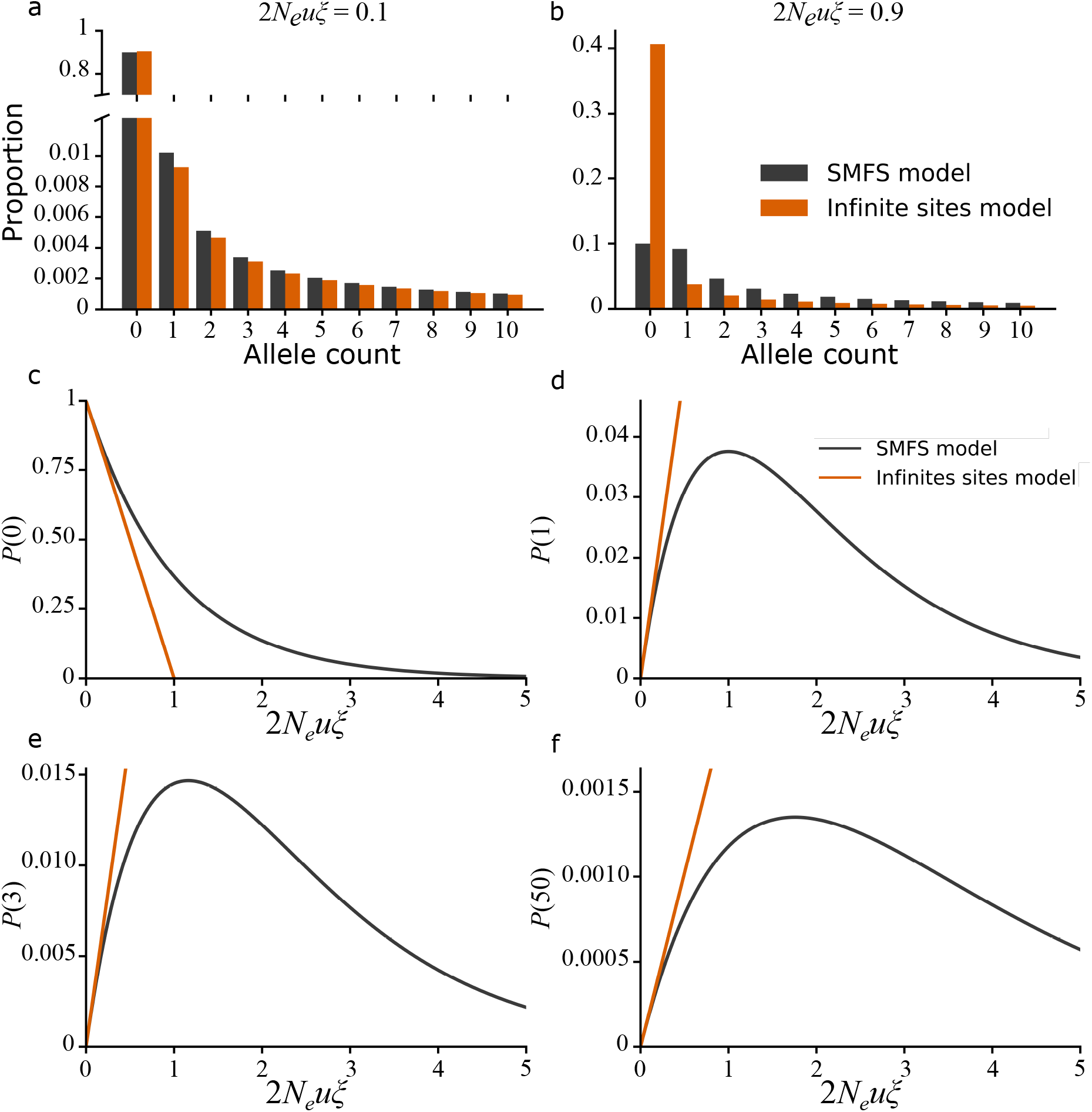
Comparison of the infinite sites model to the SMFS model. (a-b) The SFS under an infinite sites model and the SMFS model for a constant population size. The models agree when the mutation rate is low (a) but when the mutation rate is high (b), recurrent mutation changes the shape of the SFS under the SMFS model. (c-f) The proportion of sites with a given allele count as a function of the scaled mutational input 2*N*_*e*_*uξ*. (c) The proportion of non-segregating sites decays as the mutational input increase. The proportion of singletons (d), tripletons (e) and sites with an allele count of 50 (f) all increase initially but then decay when the mutational input is large.

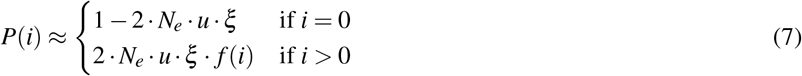

so the mutation rate only affects the proportion of segregating sites, but not the shape of the SFS for segregating sites. This is equivalent to the infinite sites approximation. For larger *u*, when 2 ·*N*_*e*_· *u*· *ξ* is of the order of 1 or larger, the shape of the SFS becomes mutation rate dependent, Fig. 2b.

For *i* = 0 (Fig. 2c), *P*(*i*) is a monotonically decreasing function of *u*, i.e. the probability of not observing the derived allele decreases as the mutational input increases. For all other allele counts (*i* > 0; Fig. 2d-f) it is non-monotonic: increasing linearly for small *u*, peaking, and then decreasing exponentially for large *u*. This reflects the fact that at low mutation rates, the probability of observing allele count *i* grows linearly as more and more sites become segregating. However, as *u* grows, more recurrent mutations are sampled, making low allele counts increasingly unlikely.

### 2.3 Relation to other models

Other models, including backward-in-time coalescent-based models^8,10^ and forward-in-time models based on branching processes^7^ or diffusion equations^38^, have included recurrent mutations.

However, at low allele frequencies the SFS under such models can be described in the form above (Eq. 6) – as a sum of convolutions of an SMFS.

For forward-in-time models^7^, if we denote the forward-in-time transition probability from an allele count of one at time *t* before present to an allele count of *i* in the present as *q*_*i*_(*t*). Then, the SFS under a forward-in-time model can be described by our formula above (Eq. 6), with

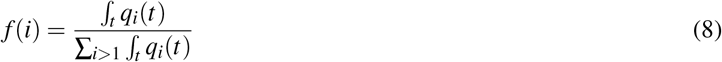

and

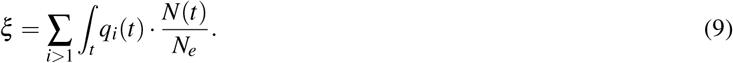

Schraiber et al^7^ calculate these transition probabilities using a branching process and arrive at results exactly equivalent to those of our model (see Supplement). These transition probabilities could also be calculated via diffusion equations^38^ or numerically via simulations^39^. Note that, unlike backward-in-time models, such forward-in-time models can readily account for selection.

For a backward-in-time model^8^, if we denote the total length of branches with *i* descendants in a gene genealogy of *m* ≫1 sampled chromosomes as *τ*_*i*_ then the SFS again takes the form of Eq. 6 with

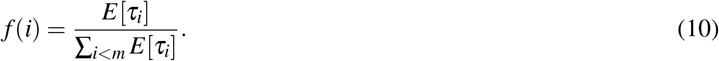

and 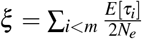. Thus, *ξ* is the expected length of all branches in a gene genealogy of sample size *m* (in units of 2*N*_*e*_). Wakeley et.al.^8^ derive these results, though they present them in a different form (see Supplement).

Our simple model can be used to extend any model that describes the frequency spectrum with a single mutational origin to describe the frequency spectrum of rare alleles with recurrent mutations. Furthermore, the use of convolutions instead of Bell polynomials^7^ or combinatorial sums^8^ vastly simplifies the formulas for, and the computation of, the SFS.

### 2.4 Challenges in predicting the SFS for differing mutation rates

Our model suggests that given the SMFS, one could predict the SFS. In this spirit, Wakeley et al.^8^ estimated the SMFS (which they describe in terms of averages over gene genealogies) from low mutation rate sites. However, there is a fundamental challenge to this approach: low mutation rate sites may still harbor recurrent mutations and will be more likely to do so as sample sizes increase, biasing estimation of the SMFS. We therefore suggest a new approach which overcomes these limitation: hierarchical estimation of the SMFS while accounting for recurrent mutation.

## 3 Combinatorial and Hierarchical Estimation of the SMFS (CHES)

There is a hierarchy in the relationship between model parameters and the observed SFS, see Fig. 3. Non-segregating sites represent zero mutational events and so their number depends only on the combination 2 ·*N*_*e*_ ·*u* ·*ξ*, an observation that serves as the basis for mutation rate estimation^10^. Singletons, i.e., sites where there is exactly one copy of the derived allele, necessarily represent a single mutational event. Thus, the number of singletons in a sample depends on 2 ·*N*_*e*_ ·*u*· *ξ*, and *f* (1). Doubletons, i.e., sites where there are exactly two copies of the derived allele, can either arise from a single mutational event or from two independent mutational events. Therefore, the number of such sites depends on 2 · *N*_*e*_ · *u* · *ξ, f* (1) and *f* (2). Similarly, *i* derived alleles can either arise from a single mutational event with *i* descendants observed in the sample or multiple mutational events each with fewer than *i* descendants observed in the sample. Therefore, the number of such sites depends on 2 ·*N*_*e*_ ·*u*· *ξ*, and *f* (1), *f* (2),…, *f* (*i*). This suggests a hierarchical inference approach: We first infer *ξ* from the mutation rates of segregating versus non-segregating sites. We then infer *f* (1) from *ξ* and the number of singletons. Next, we infer *f* (2) from *ξ, f* (1) and the number of doubletons. We continue this process iteratively for higher and higher values of *i*. We next provide mathematical details for this process, which we name Combinatorial and Hierarchical Estimation of the SMFS, or CHES:

**Figure 3.**
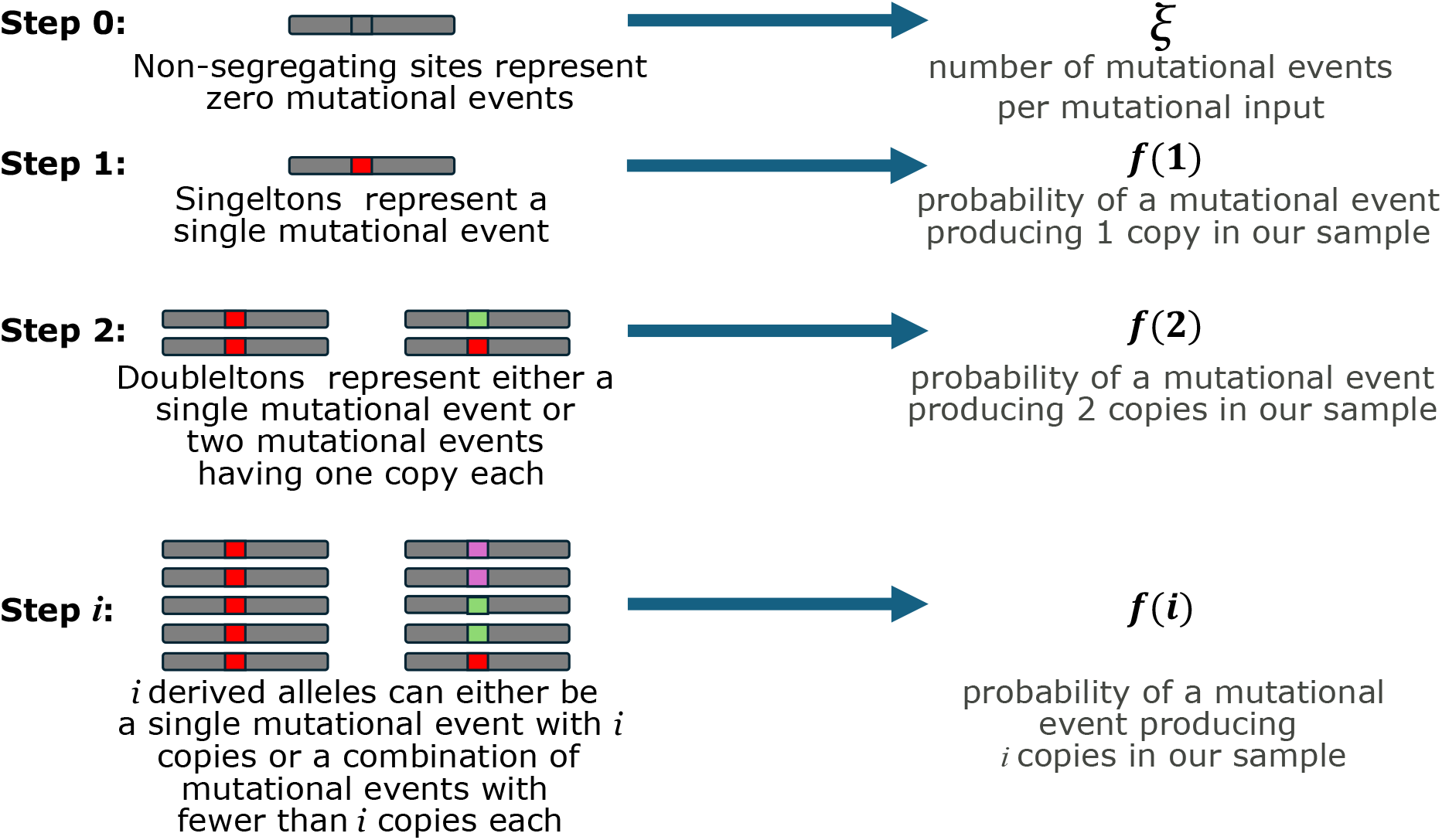
Combinatorial hierarchical estimation of the SMFS (CHES) overview.

### Step 0: Estimating the number of sampled mutational events per mutational input (*ξ*)

We first infer the number of sampled mutational events per mutational input (*ξ*) from the distribution of segregating sites. We treat site segregation as a Bernoulli variable, *x*_*l*_, for each site *l* with:

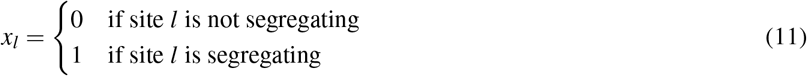

Since a non-segregating site has zero mutation events, *P*(*x*_*l*_ = 0) = *P*(*i* = 0) = *P*(*k* = 0 | *N*_*e*_, *u*_*l*_, *ξ*), and the expected total number of non-segregating sites is *N*(0) = ∑_*l*_ *P*(*k* = 0 | *N*_*e*_, *u*_*l*_, *ξ*). Therefore, the likelihood function for *ξ* is given by,

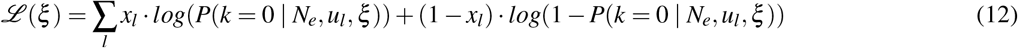

which we then maximize to obtain an estimate of 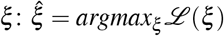.

We assume that the mutation rates, *u*_*l*_, have already been estimated. In the supplement, we detail how this could be extended to include uncertainty in the estimation of *u*_*l*_, i.e a distribution of possible mutation rates for each site. When applying this method to non-human data, it might be necessary to first estimate the mutation rates, see Seplyarskiy et al.^10^.

### Step 1: Estimating *f* (1) from singletons

A site with a single derived allele can only represent a single mutational event. Therefore, the probability that a site is a singleton is 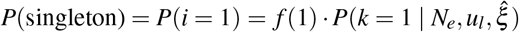, and therefore the expected total number of singletons is:

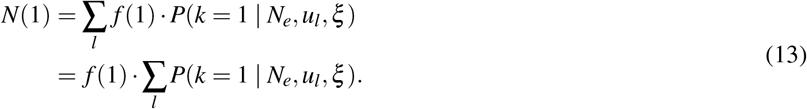

suggesting an estimator for *f* (1),

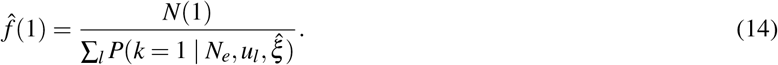

### Step 2: Inferring *f* (2) from doubletons

A site with two derived alleles can represent either a single unique mutation with two copies or two unique mutations with one copy each. That is, *P*(doubleton) = *P*(*i* = 2) = *f* (2) · *P*(*k* = 1 | *N*_*e*_, *u*_*l*_, *ξ*) + (*f* (1))^2^ · *P*(*k* = 2 | *N*_*e*_, *u*_*l*_, *ξ*) and therefore the expected total number of doubletons is:

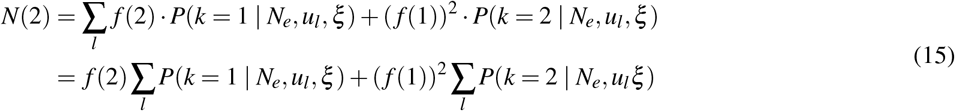

Therefore, we can estimate *f* (2) using 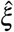 and 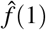 by:

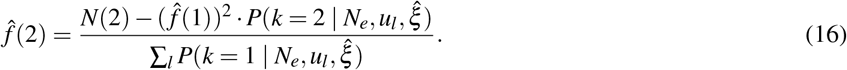

### Step *i*: Inferring *f* (*i*) from SFS

We can now generalize this process. The probability of seeing *i* derived allele copies is: *P*(*i*) = ∑_*k*_ *f* ^\**k*^(*i*) *P*(*k*|*N*_*e*_, *u*_*l*_, *ξ*). By definition, *f* ^\**k*^(*i*) depends only on values of *f* (*j*) for which *j*≤*i* −*k*, and therefore, having estimated *f* (*j*) for *j* < *i* we now have estimates for 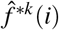 for all *k* except *k* = 1. We therefore split the above expression into:

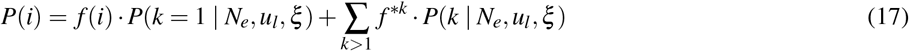

and therefore:

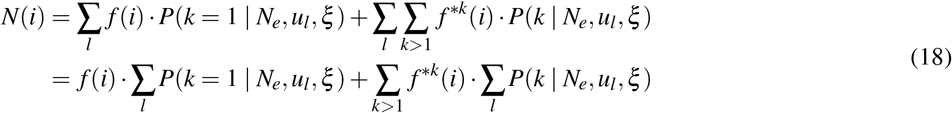

This suggests we can estimate *f* (*i*) by:

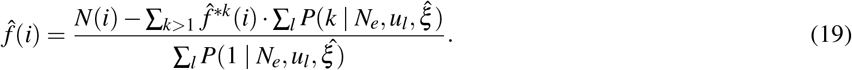

That is, we subtract from the observed number of sites with allele count *i* the expected number of sites with allele count *i* that are the result of recurrent mutation to estimate the number of site with allele count *i* due to a single mutation. We then divide this number by the expect number of sites with a single mutation to get the proportion of sites with a single mutation and allele count *i*, that is *f* (*i*).

## 4 Results

### 4.1 Evaluating hierarchical estimation of the SFS with simulations

To evaluate the ability of CHES to recover the true SMFS, we conducted forward-in-time simulations in an exponentially growing population. We simulated a diploid population growing from a population size of 10^4^ to a population size of 10^8^ in 1, 000 generations. In the simulations, the dynamics of rare alleles are specified by a Poisson branching process. Mutation rate distributions are drawn from estimated distributions for human synonymous substitutions (Schraiber et al.^36^; see Supplement) and the number of simulated sites is chosen to match the number seen in human whole exome sequencing. When we look at the allele count distribution for derived alleles, we see a clear dependency on mutation rate, as expected (Fig. 4B-C).

**Figure 4.**
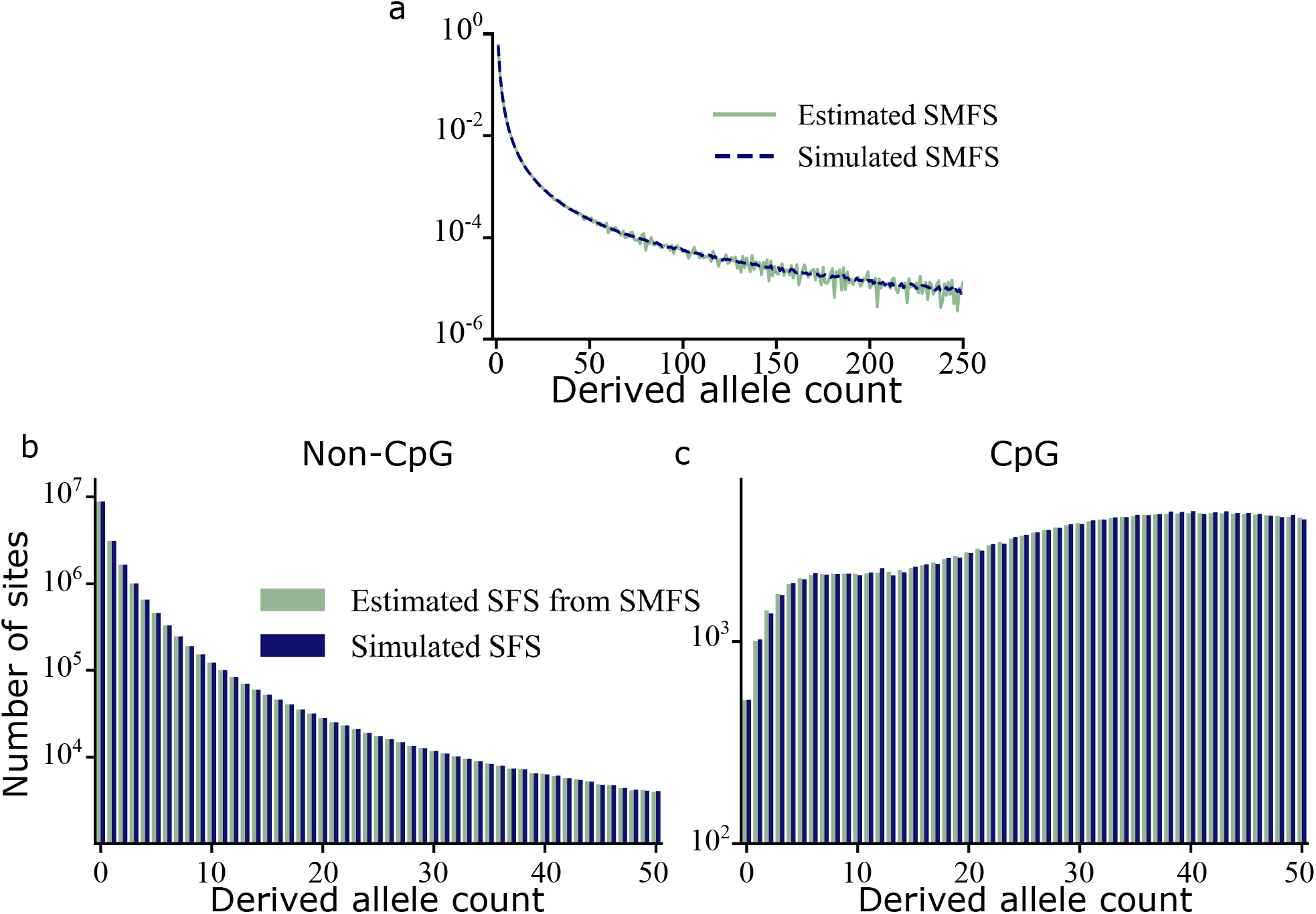
Applying CHES to simulated data. (a) Estimated SMFS from CHES and true simulated SMFS from IBD alleles in simulated data. This SMFS was used to produce the predicted SFS in b & c. (b-c) The simulated SFS for non-CpG and CpG mutations and the SMFS-based predictions of this SFS.

When applied to simulations, CHES accurately recovers the true SMFS, up to an allele count of *i* = 200 (Fig. 4). Beyond *i* = 200, the number of sites with a given allele count becomes low (of the order of 10 or less). Since the SFS is noisy then, because of the hirarchical nature of CHES, the noise accumulates and the SMFS beyond *i* = 200 becomes dominated by noise. The inferred SMFS can recapture the SFS both at low mutation rate (Fig. 4B) and high mutation rate (Fig. 4C) sites.

When applying CHES to simulations, we can use the true mutation rate at each site. However, in real data, the mutation rates are unknown. Thus, in practice, we assume that the mutation rates follow the distribution of mutation rates estimated by Schraiber et al.^7^. To test the reduction in power due to this limitation, we ran CHES on our simulations using the true mutation rate at each site. We observed essentially identical results with no power gain (see Supplement) suggesting that estimates of the distributions of mutations rates are sufficient to infer the SMFS.

### 4.2 Human SFS

We applied CHES to an exome sequencing dataset binned by trinucleotide context and methylation level (Fig. 5A, and see^7^). The inferred SMFS is very similar to the SFS of low mutation rate sites because at this sample size, low mutation rate sites rarely exhibit recurrent mutation, 5B. However, CHES produces a much less noisy estimate of the SMFS than simply using sites with sufficiently low mutation rates (as used in Wakeley et al.^8^), since such sites are rare. An alternate version of CHES which accounts for sequencing errors produced near-identical results because of the low error rate (see supplement), so we used the simpler method for all tests reported here.

**Figure 5.**
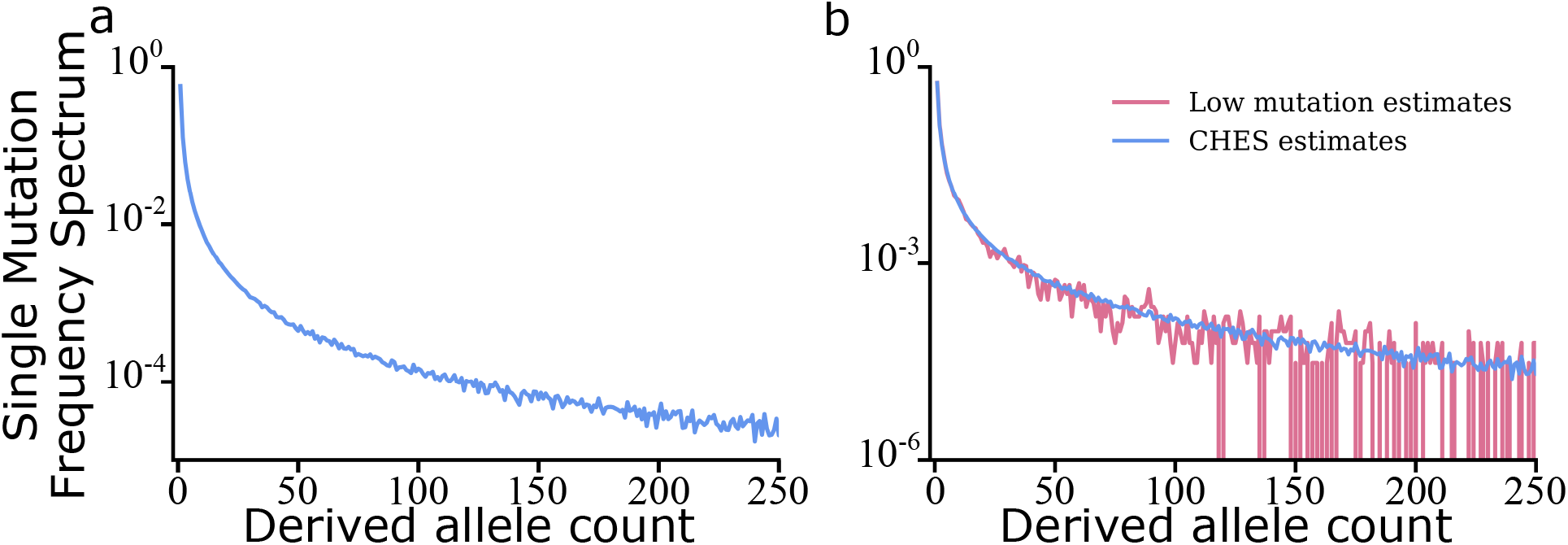
The inferred single mutation frequency spectrum (SMFS) for synonymous mutations in 1,000,000 exomes. (a) The CHES-inferred SMFS. (b) The SFS for mutations with low mutation rate (scaled by the mutational input 2*N*_*e*_*u*) provides a noisy approximation for the SMFS.

The shape of the SMFS strongly deviates from that predicted by a neutral, constant population size model, where the SMFS scales inversely to allele frequency 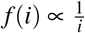, Fig. 6. However, the SMFS also deviates from the predictions of an exponential population growth model, where the SMFS scales approximately with 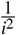 ^8,40^. Instead, the SMFS seems to follow an intermediate scaling with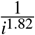. This finding is puzzling, since both census population data and population genetic data suggest that recent population growth has been super-exponential^15,41^. We will address the question of how super-exponential growth can lead to an SMFS intermediate between constant and exponentially growing population models in an upcoming publication.

**Figure 6.**
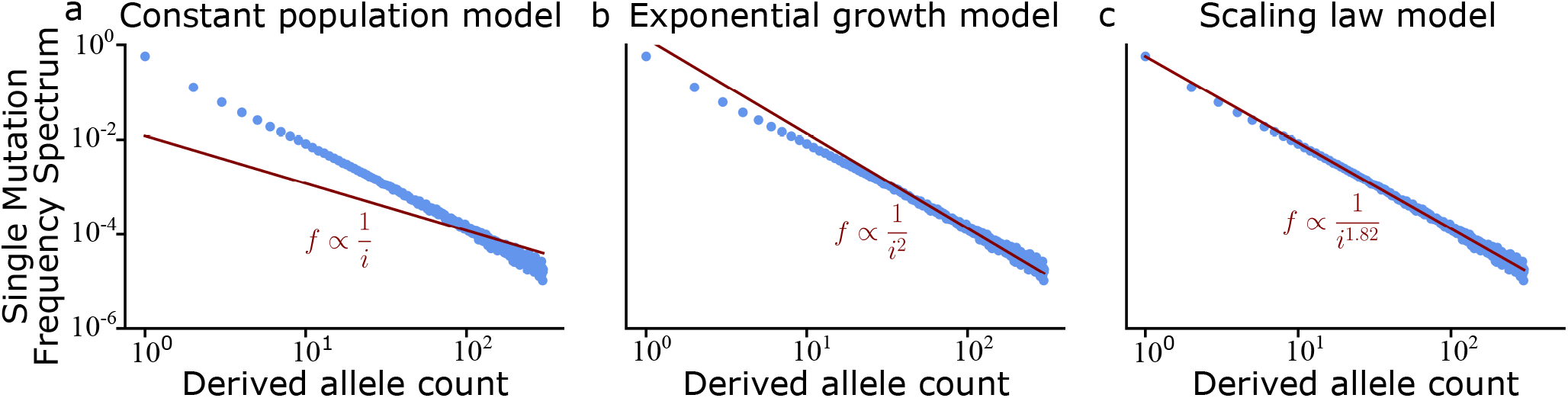
Fitting a scaling law to the SMFS. (a) A constant population size scaling law. (b) A scaling law for exponentially growing populations. (c) The general scaling law that best approximates the SMFS of the human exome dataset.

With the inferred SMFS from CHES, we see that differences in mutation rates explain the difference in SFS between synonymous methylated CpG mutations (with high mutation rate) and all other synonymous mutations, Fig. 7.

**Figure 7.**
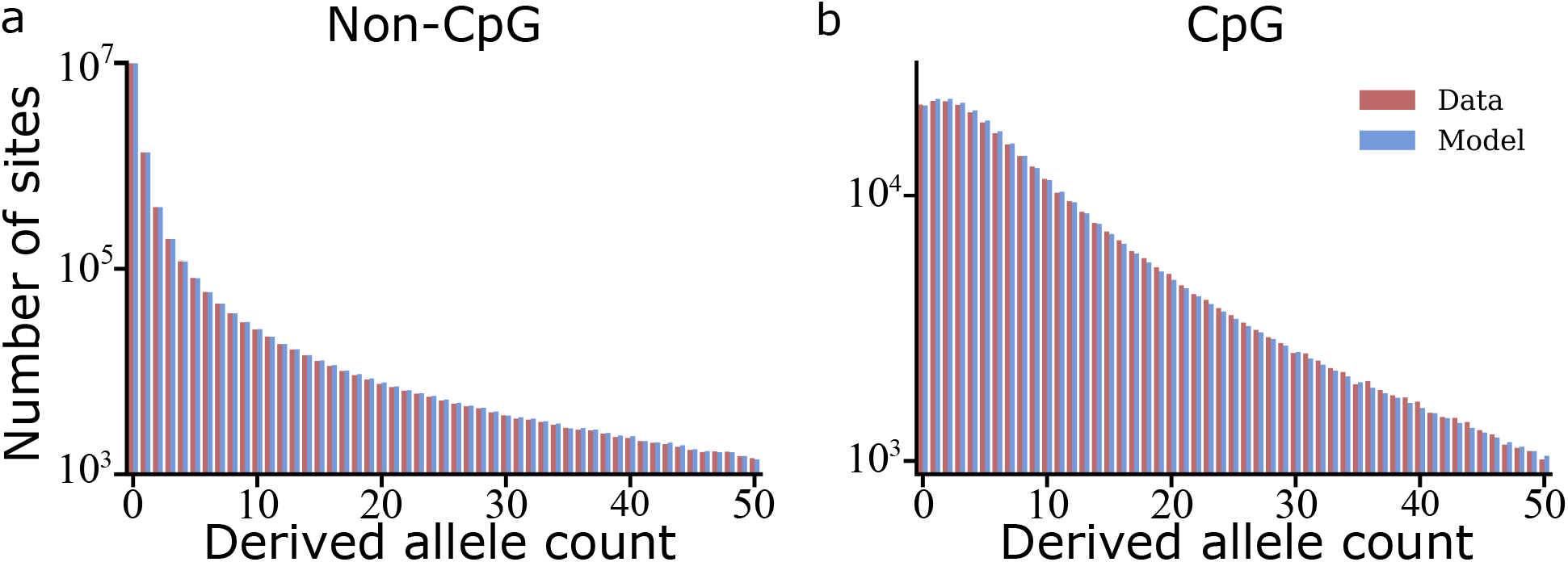
Difference in the site frequency spectrum (SFS) between (a) non-CpG and (b) methylated CpG C to T muatations are well explained by the CHES-inferred SMFS and differences in mutation rate.

### 4.3 Signal of selection

The inferred SMFS allows us to test for deviation from neutrality at selected sites. We repeated Schraiber et al.’s data analysis pipeline^7^ to produce an SFS for missense and stop-gain mutations binned by their trinucleotide context and methylation level. If we compare the predictions of our model with the inferred SMFS, we see a pronounced deviation from the neutral expectation produced by the synonymous SMFS. This deviation is stronger at stop-gain than missense mutations, Fig. 8.

**Figure 8.**
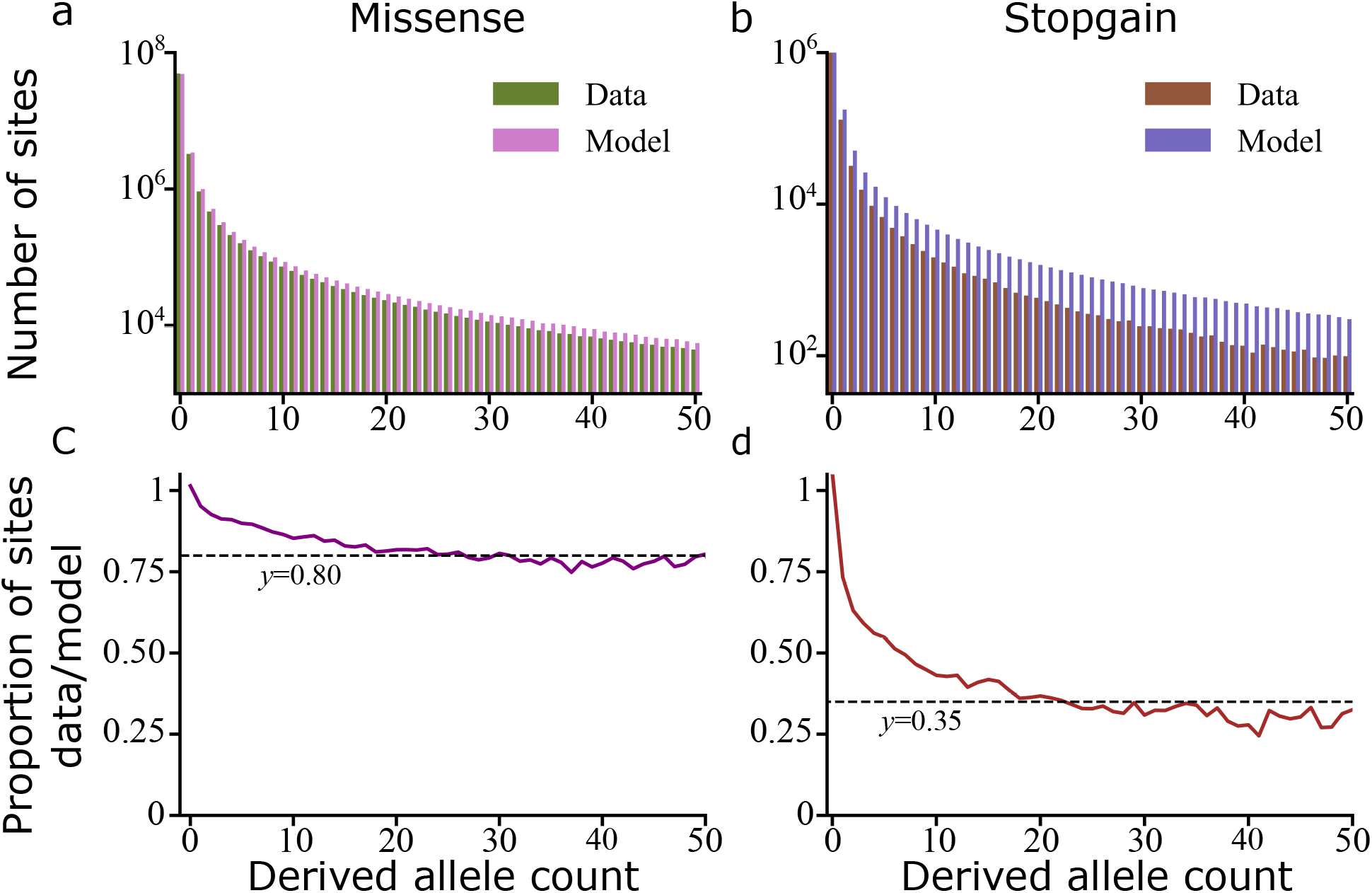
We compare the prediction of our SMFS model, with parameters inferred by CHES, to the SFS of (a) missense and (b) stop-gain mutations. Fewer missense and stop-gain mutations have high allele counts. The near-constant ratio between the observed and predicted number of mutations (c-d) for allele counts above 10 indicates that ~ 20% of missense and ~ 65% of stop-gain mutations are under strong selection.

We can interpret these deviations by looking at the ratio of the observed to the expected SFS, Fig. 8c-d. Except for some deviations at low allele counts, this ratio is approximately constant. This indicates that we observe fewer segregating variants at appreciable allele counts (AC>10) than expected, but their SFS is similar in shape to the expected SFS. We interpret the observed signal as coming from effectively neutral mutations while the reduction is due to the removal of deleterious mutations by selection pressure. Simulations show that for such low allele frequencies, only highly deleterious variation (with selection coefficient > 1%) would be censored by selection. We therefore interpret our results as indicating that 20% of missense mutations in the exome are strongly deleterious, while 65% of stop-gain mutations are strongly deleterious.

These results are consistent with previous estimates of selection strength on coding regions^26^ and stop-gain mutations^27^, though those estimates used smaller datasets and therefore did not need to account for recurrent mutation. While this method to estimate selection is similar in spirit to existing measures of selective constraint (Mutation Adjusted Proportion of Singletons (MAPS)^5^ and its extension CAPS^28^), it uses the full frequency spectrum, accounts for recurrent mutation, and produces a value — one minus the observed-to-expected SFS ratio — that can be directly interpreted as the proportion of mutations under strong selection.

## 5 Discussion

In this work, we introduce a new approach for modeling the frequency distribution of rare alleles with recurrent mutation. Unlike previous approaches, we define a single mutation frequency spectrum (SMFS) which is the frequency spectrum of identical-by-descent rare alleles. The overall frequency spectrum is then a mutation-rate-dependent weighted sum of convolutions of the SMFS. This approach separates the population genetic processes giving rise to the SMFS, such as demography and selection, from mutation recurrence. We show how this approach provides a unified framework for previous results from both forward-in-time and backward-in-time models. After developing a model for how the SFS depends on the SMFS and mutation rate, we develop a procedure, CHES, to estimate the SMFS from the SFS and estimated mutation rate.

We applied CHES to a human exome-sequencing dataset^5–7^ to show that variation in SFS between low and high mutation rate sites is well-explained by our model. We further show that the shape of the SMFS is intermediate between a constant population size and an exponential population growth model; given the abundant evidence for recent super-exponential growth in human populations, this observation suggests the need for further work to understand how demographic history shapes the SMFS. Lastly, we used our inferred model to detect the effects of selection on the frequency spectrum of missense and stop-gain mutations.

Our model can serve as an intermediary between the mutation-rate-independent predictions of the infinite sites model and empirical results emerging from genomic data. It provides an intuitive and computationally simple way to model the SFS given the results of an infinite sites model, and therefore an easy way to infer the parameters of such a model from data which includes recurrent mutations. Furthermore, one could apply existing methods that do not account for recurrent mutation directly to the SMFS. For example, one could combine the CHES estimated SMFS with Poisson Random Field methods to estimate demographic history and the strength of natural selection^11–15,42^. Furthermore, while we used our model to qualitatively estimate the strength of selection on missense and stop-gain mutations, there is clearly room to develop more elaborate SMFS-based methods that will provide a quantitative estimate of the strength of natural selection on such mutations. Our estimation of the SMFS is fast and simple, but it is far from optimal. Our inference relies solely on combinatorics and algebra to infer the SMFS, and its hierarchical nature means it infers the SMFS entries one by one. It is clear to us that more sophisticated methods can be developed and would probably prove useful as sample sizes increase, since CHES relies on there being many sites where only a single mutational event has been sampled, i.e. where all derived alleles are IBD. Furthermore, such methods may be necessary to provide insight into the distribution of selection coefficients (known as the DFE) at functional sites, since CHES assumes that all sites have the same SMFS and this assumptions is violated when the SMFS depends on the selection coefficient.

A completely different approach would be to estimate the number of independent mutational events at a site and estimate the number of sampled allele copies descendant from each such event^43,44^. This could be achieved by looking at the haplotypic structure around the site. Given a low enough recombination rate and enough common variation around the focal site, alleles from different mutational events should have distinct genetic backgrounds. A phylogenetic analysis (e.g., ancestral recombination graph reconstruction) for the rare alleles at a site should reveal distinct clades, each corresponding to a different mutational event, which would allow for a direct estimation of *ξ* and *f* (*i*). One such approach^44^ that concentrated only on doubletons successfully distnguished between IBD and non-IBD doubletons in African *Anopheles gambiae* samples. Another^43^, more general, method was applied to the UK10K dataset^45^, which consists of 3,621 individuals. At this modest sample size, this method could detect some instances of non-IBD alleles, but its extension to larger datasets with many mutational origins at many sites remains untested.

The growing size of genomic datasets provides many new theoretical, computational, and conceptual challenges to population genetics, which we hope this work will provide a stepping stone for addressing.

## Data and code availability

Data can be found at 10.6084/m9.figshare.32444397

Code can be found at https://github.com/Deepgh/smfs-inference

## Acknowledgements

The authors wish to thank John Wakeley, Louis Fan, Maryn Carlson, Evan Koch, Arbel Harpak, Jeffery Spence, Carl Veller, Pavitra Muralidhar, and John Novembre for helpful discussions.

